# Comprehensive characterization of mitochondrial bioenergetics at different larval stages reveals novel insights about the developmental metabolism of *Caenorhabditis elegans*

**DOI:** 10.1101/2024.06.26.600841

**Authors:** Danielle F. Mello, Luiza Perez, Christina M. Bergemann, Katherine S. Morton, Ian T. Ryde, Joel N. Meyer

## Abstract

Mitochondrial bioenergetic processes are fundamental to development, stress responses, and health. *Caenorhabditis elegans* is widely used to study developmental biology, mitochondrial disease, and mitochondrial toxicity. Oxidative phosphorylation generally increases during development in many species, and genetic and environmental factors may alter this normal trajectory. Altered mitochondrial function during development can lead to both drastic, short-term responses including arrested development and death, and subtle consequences that may persist throughout life and into subsequent generations. Understanding normal and altered developmental mitochondrial biology in *C. elegans* is currently constrained by incomplete and conflicting reports on how mitochondrial bioenergetic parameters change during development in this species. We used a Seahorse XFe24 Extracellular Flux (XF) Analyzer to carry out a comprehensive analysis of mitochondrial and non-mitochondrial oxygen consumption rates (OCR) throughout larval development in *C. elegans*. We optimized and describe conditions for analysis of basal OCR, basal mitochondrial OCR, ATP-linked OCR, spare and maximal respiratory capacity, proton leak, and non-mitochondrial OCR. A key consideration is normalization, and we present and discuss results as normalized per individual worm, protein content, worm volume, mitochondrial DNA (mtDNA) count, nuclear DNA (ncDNA) count, and mtDNA:ncDNA ratio. Which normalization process is best depends on the question being asked, and differences in normalization explain some of the discrepancies in previously reported developmental changes in OCR in *C. elegans*. Broadly, when normalized to worm number, our results agree with previous reports in showing dramatic increases in OCR throughout development. However, when normalized to total protein, worm volume, or ncDNA or mtDNA count, after a significant 2-3-fold increase from L1 to L2 stages, we found small or no changes in most OCR parameters from the L2 to the L4 stage, other than a marginal increase at L3 in spare and maximal respiratory capacity. Overall, our results indicate an earlier cellular shift to oxidative metabolism than suggested in most previous literature.

## 1. Introduction

Mitochondria play key roles in development and disease [1–3]. Understanding how genetic and environmental factors impact mitochondrial function is critical, because ∼1 person in 4,000 suffers from a mitochondrial disease [4], and many pollutants [5] and drugs [6] cause mitochondrial toxicity. Mitochondria are also key mediators of cellular stress and immune responses [7].

The model organism *Caenorhabditis elegans* has been used extensively to study developmental biology and genetics [8], mitochondrial disease [9], and mitochondrial toxicity [10]. *C. elegans* development includes the egg stage (in which ∼50% of the final number of somatic cells are made), followed by 4 larval stages and adulthood. Early work by Lemire and colleagues demonstrated that pharmacologically inhibiting mitochondrial function during development blocked or slowed developmental progression from the L3 to the L4 larval stages [11], as did blocking mitochondrial DNA (mtDNA) replication pharmacologically [12]. Strains carrying mutations in genes encoding mitochondrial proteins [11] or the DNA polymerase responsible for mtDNA replication of [13] also showed slowed or blocked development, as did worms exposed to a variety of pollutants that targeted mitochondria [14, 15]. Lower levels of mitochondrial stress resulting from RNA inhibition that still permit development to adulthood can lead to both deleterious and beneficial outcomes later in life [16–18]. Similarly, both deleterious and beneficial outcomes later in life can result from lower levels of mitochondrial stress induced by chemical exposures during development [19, 20]. Mitochondrial toxicity also affects *C. elegans* neuronal function [21, 22], immune function [23, 24], and other cellular processes [25–27] impacted by mitochondrial dysfunction in higher eukaryotes. However, while *C. elegans* is generally an excellent model for studying how genetic and environmental factors affect mitochondrial function during development, this utility is limited by incomplete and in some cases conflicting literature reports of how mitochondrial biology changes during development in this organism.

The integrity and transcription of the mitochondrial genome is necessary for aerobic energy production, because mtDNA encodes 12 (in worms) or 13 (in humans) core proteins of the mitochondrial respiratory chain (MRC). The mtDNA copy number (mtDNA CN, or number of mitochondrial genomes per cell) is highly variable by cell type, and undergoes a developmental bottleneck because mtDNA is not replicated in early stages after oocyte fertilization, but rather simply allocated into new cells, with replication beginning around the blastocyst stage in vertebrate species examined [28]. In *C. elegans*, Tsang and Lemire first showed that mtDNA CN increased substantially in development [12]. They also showed that when mtDNA replication was halted using the mtDNA-selective replication inhibitor ethidium bromide, nematodes did not develop past L3 [12]. Trifunovic and colleagues [13] later showed that loss of the sole replicative DNA polymerase also blocked development to adulthood, and Leung, Bess and colleagues found that mtDNA damage slowed larval development [14, 29, 30]. The fact that mtDNA CN either decreases slightly [12], stays constant [13] or increases only slightly [29] from the embryo to the L1 stage shows that a significant bottleneck must also occur in this species, since nuclear DNA (ncDNA) CN increases ∼500-fold during this time. All these reports described a general increase in mtDNA content and requirement for mtDNA function during development, with quantitative discrepancies that might be explained by methodological differences.

Similarly, in general, mitochondrial bioenergetics and metabolism change throughout development [31]. Characterizations of *C. elegans* aerobic metabolism during larval development have yielded somewhat conflicting results. Vanfleteren and De Vreese [32] reported peak protein-normalized OCR at the L3 and L4 stages, but Houthoofd et al. reported peak basal respiration at the L2 stage, after normalization to protein content [33]. Huang and Lin [34] developed a novel microfluidic device that permitted measurement of OCR on individual nematodes during development except for the L1 stage, at which OCR was not detectable with their methodology. Using this device, they reported steadily increasing OCR from L1 to L4 stages, as measured on a per-worm basis. They were also able to measure some specific aspects of OCR (e.g., ATP-linked, non-mitochondrial OCR, proton leak, and spare respiratory capacity), but reported these only at the L3, L4, and adult stages. They highlighted low ATP-linked OCR (on a per-worm normalization basis) at the L3 stage, and suggested that this corresponded to largely glycolytic energy production. Overall, the existing literature offers some insight into how mitochondrial bioenergetics change through *C. elegans* development. However, discrepancies exist, and a full time-course normalized to mitochondrial amount and incorporating analysis of the different components of oxygen consumption is lacking.

The goal of this study was to provide a comprehensive description of mitochondrial and non-mitochondrial OCR throughout larval development in *C. elegans*. We used a Seahorse XFe24 Extracellular Flux Analyzer to analyze whole animal basal OCR, basal mitochondrial OCR, ATP-linked OCR, spare respiratory capacity, and non-mitochondrial OCR, after optimizing the conditions for each of these at each larval stage. There are many ways in which OCR data can be normalized, each with its own strengths and limitations; we present and discuss results as normalized per individual worm, protein content, worm volume, mtDNA CN, ncDNA CN, and mtDNA:ncDNA ratio. Our results demonstrate when normalized to total protein, worm volume, or ncDNA or mtDNA count, after a 2-3-fold increase from L1 to L2 stages, changes in most OCR parameters from the L2 to the L4 stage are relatively modest. This indicates an earlier cellular shift to oxidative metabolism than suggested in most previous literature.

## 2. Methods

### 2.1 Synchronization of Nematodes

Worms were prepared using the hypochlorite bleach method of egg preparation [35]. A plate of mixed-stage worms was washed with K-medium (50 mM NaCl, 30 mM KCl, 10 mM NaOAc; pH 5.5) (K-med) solution and transferred to a 15mL tube. The worms were then washed twice with 15 mL of K-medium, and gravid adult worms were allowed to settle by gravity. We isolated the adults by aspirating the supernatant, then isolated eggs by treating gravid adults with hypochlorite bleach (1 mL 5N hypochlorite bleach, 500 µL 1M NaOH, and 3.5 mL K-medium). The 15mL tube containing the adult worms in bleach was placed in a 20 °C incubator on a shaker for 8 minutes. We briefly removed tubes to vortex and observed changes under the microscope every two minutes. When only fragments of worms could be seen, we increased the volume to 15mL with K-medium solution to stop the reaction and washed the eggs with K-medium twice, centrifuging between washes. Next, we placed 10µL of the egg solution in three drops on a slide and counted the number of eggs in 10 µL. We transferred eggs to OP50-seeded K-agar plates [36] supplemented with 6.75 µM (final in-agar concentration) nystatin and incubated at 20°C to allow development. We plated about 1000 eggs per plate for experiments with L1s; about 500 for experiments with L2s and L3s; and about 300 for experiments with L4s to avoid food deprivation during development. We removed plates from the incubator when worms reached the desired larval stage, which corresponded to 21h for mid-L1s, 34h for mid-L2s, 43h for mid-L3s, 46h for late-L3s, and 53h for mid-L4s.

### 2.2 Seahorse XFe24 Extracellular Flux Analyzer-based measurements of oxygen consumption rate

#### 2.2.1 Preparation and counting of nematodes

Upon reaching the desired larval stage, we harvested nematodes by washing the OP50-seeded K-agar plates with K-medium into 15mL centrifuge tubes. Tubes were centrifuged for 30 seconds at 2200 RCF if worms were L1-L3s; we let L4s settle by gravity. We resuspended worms in the centrifuged tubes with K-medium and placed them in an orbital shaker for 20 minutes at 20°C to allow gut clearing. Tubes were then centrifuged for 30 seconds at 2200 RCF (regardless of larval stage) and re-suspended in K-medium. To estimate the number of worms per microliter, we suspended the worms and transferred 3-4 drops of 20µL of worms to a glass slide and counted under a microscope. We calculated the concentration of worms per microliter and the volume needed to achieve the desired concentration depending on the nematode’s development stage, given 525 µL total volume added to each well. The desired number of worms (we tested 1000-3000 for L1s; 300-700 for L2s; 150-250 for L3s; and 75 for L4s; the ideal number yielding OCR rates of 200-400 pmol oxygen/minute for the basal reading had been previously optimized for the L4 stage but not earlier stages) was transferred to the 24-well Seahorse plate, and if needed the wells were completed with K-medium to reach a total volume of 525 µL.

#### 2.2.2 Preparation of Seahorse XFe24 Extracellular Flux Analyzer and assay plates

We hydrated the Seahorse sensor cartridge overnight using 1mL per well of the XF Calibrant solution at room temperature. The Seahorse software was set up by identifying injection strategies along with final ETC inhibitor concentration, solvent and percent solvent used in each port. Configurations were set to include 8 cycles of Basal Measurements, 1 mix cycle to oxygenate the micro-chamber, 3 wait cycles to allow the worms to settle, and a measuring cycle to 3 minutes [37, 38]. 8 measurement cycles were used for measures of FCCP response, 14 for DCCD response and 4 for sodium azide response. 75 µL of the desired concentrations of each ETC-inhibitor were transferred to the appropriate port (FCCP or DCCD to Port A and sodium azide to Port B, both dissolved in DMSO and stored at -80 C until use as described [38]). Once appropriate ETC inhibitors were added to injection ports, the cartridge was loaded onto the Seahorse XFe24 Extracellular Flux Analyzer for calibration. Upon calibration, the utility plate (with XF Calibrant solution) was replaced with a 24-well plate containing nematodes for the OCR measurement assay to begin. The data collection process took about three hours per plate.

#### 2.2.3 OCR Analysis

After each equipment OCR run, raw graphs were analyzed and wells that presented serious technical problems were discarded from the calculations. Problems that resulted in exclusion were: drug injection did not cause any changes in OCR; negative OCR levels; non-linear oxygen consumption caused by oxygen depletion (which can be seen by analyzing the raw oxygen levels). Equations and measurement timepoints used to calculate each bioenergetic parameter are shown in Table 1.

| Parameter | Equation |
| --- | --- |
| **Non-mitochondrial OCR** | Average rate of 3 last measurements after sodium azide injection |
| **Total Basal OCR** | Average rate of all measurements except the first and the last |
| **Mitochondrial Basal OCR** | $(\text{Total Basal OCR}) - (\text{Non-Mitochondrial OCR})$ |
| **Maximal OCR** | 3 highest rate measurements after FCCP injection |
| **Spare capacity** | $(\text{Maximal OCR}) - (\text{Total Basal OCR})$ |
| **ATP-linked OCR** | $(\text{Total Basal OCR}) - (2 \text{ lowest rate measurements after DCCD injection})$ |
| **Proton leak** | $(\text{ATP-linked OCR}) - (\text{Non-Mitochondrial OCR})$ |

### 2.3 Mitochondrial and nuclear DNA copy number

We measured mtDNA and ncDNA copy numbers per nematode as described [39]. We transferred 6 individual worms using the platinum worm pick into PCR tubes containing 90µL of 1x worm lysis buffer and froze them in the -80°C freezer. After at least 10 minutes, we thawed the samples, vortexed and spun them briefly. We placed them in a thermal cycler to heat to 65 °C for 1 hour, then 95°C for 15 minutes, then hold at 8 °C. The worm lysate was used as template DNA. We prepared a standard curve by thawing an aliquot of mtDNA copy number standard curve plasmid (10,000,000 copies/µL) and diluted it to 32,000 copies/µL. We serially diluted it to 4,000 copies/µL. The standard curve contains 64,000, 32,000, 16,000, 8,000, 4,000 and 0 copies per well. We then added 2µL of the worm lysate, 2µL of the mtDNA specific primer pair (400nM), 12.5µL SYBR Green PCR Master Mix), and 8.5µL of water in each well of the 96-well PCR plate. We amplified the target DNA using the ABI 7300 Real Time PCR system at 50 °C for 2 minutes, 95°C for 10 minutes, 40 cycles of 95°C for 15 seconds. We selected the option to calculate a dissociation curve for each sample to ensure the presence of a single product. The ncDNA copy number was quantified using the same method, however using a nuclear DNA-specific primer. The standard curve was prepared using the lysate of the *glp-1* mutant *C. elegans* which does not have a germline and 40µL of nuclease-free water (784 copies/µL). We serially diluted 1:1 the lysate until reaching 24.5 copies/µL.

### 2.4 Total protein content

Aliquots (2-3 replicates per experiment) of worms (∼2000 for L1s and L2s; ∼1000 for L3s; and ∼500 for L4s) obtained from same sample used for the OCR measurements were transferred to 1.5mL tubes and frozen at -80°C for total protein extraction. Once thawed, they were centrifuged, and the supernatant was removed. 150 µL of 10% SDS was added to the nematode sample, and the samples were ultrasonicated (Model 3000 Ultrasonic Homogenizer, BioLogics, Inc.) for 2 cycles of 30s with the lowest amplitude setting. Complete nematode lysis was confirmed under the stereoscope. Protein content was measured using the Pierce™ BCA Protein Assay Kit (Thermo Scientific). Standard curve contained known concentrations of bovine serum albumin (1000, 750, 500, 250, 125, 125, 0 ug/ml) in the presence of 10% SDS.

### 2.5 Volume analysis

An aliquot of worms (100-200 individuals) obtained from same sample used for the OCR measurements was transferred to 6 cm K-agar plates lacking Bacto Peptone and allowed to air dry at 20 °C. Images were taken through automated scanning of the plate using the Keyence BZ-X700 microscope. Images were analyzed using the Worm Sizer plugin for ImageJ/Fiji [40]. Using this plugin, to minimize variation in measurement of worm volume, it is important to keep the image settings consistent between experiments, because worm volume is influenced by shadows cast onto the plate. This effect can be minimized by increasing the aperture stop.

### 2.6 Statistical analyses and graphical presentation

One-way ANOVA followed by Tukey’s multiple comparisons test were applied using the ANOVA function in the car package in R version 4.2.1. Each experiment was repeated 2-4 times, with replicate wells on the same plate (which were aliquoted from the same group of cultured worms, which was counted as an independent biological replicate) counted as technical replicates. Error bars in graphs indicate standard errors of the mean, except in cases where n = 2, in which cases the bars indicate the two data points.

### 2.7 Data reporting

All data are available in **File S1**.

## 3. Results

### 3.1 Optimized conditions for OCR measurements in all larval stages

To establish the profile of *C. elegans* mitochondrial bioenergetics throughout different life stages we stage-synchronized wild-type (N2) nematodes (**Figure 1A**) and measured OCR levels with and without different drugs (electron transport chain inhibitors and a mitochondrial uncoupler). Optimization of worm number per well is critical to ensure that enough worms are present for reliable measurements (well above the background OCR and detection limits), but not so high that oxygen is depleted to the point of becoming limiting for cellular processes or for the reliability of the Seahorse Xfe24 Extracellular Flux Analyzer data (e.g. so high that re-oxygenation step is impaired). Drug optimization is critical both to ensure that a full response is observed, and to avoid a toxic response resulting in misleadingly decreased OCR. For example, insufficient uncoupling with FCCP will yield a falsely low estimate of maximal OCR and spare respiratory capacity. Too much uncoupling, however, will kill cells (and worms), leading to a decrease in OCR. Substrate depletion-mediated decreases in OCR may also be observed at later timepoints after FCCP injection; these timepoints should be excluded (although differences in time to depletion may be informative of substrate availability). Therefore, it is critical to optimize FCCP concentrations to show a maximal but steady OCR plateau but not a peak followed by a sharp decrease in OCR. We previously [16, 37, 41] reported optimization of number of worms per well and drug concentrations for a subset of larval stages, and here report optimized parameters for all larval stages.

**Figure 1.**
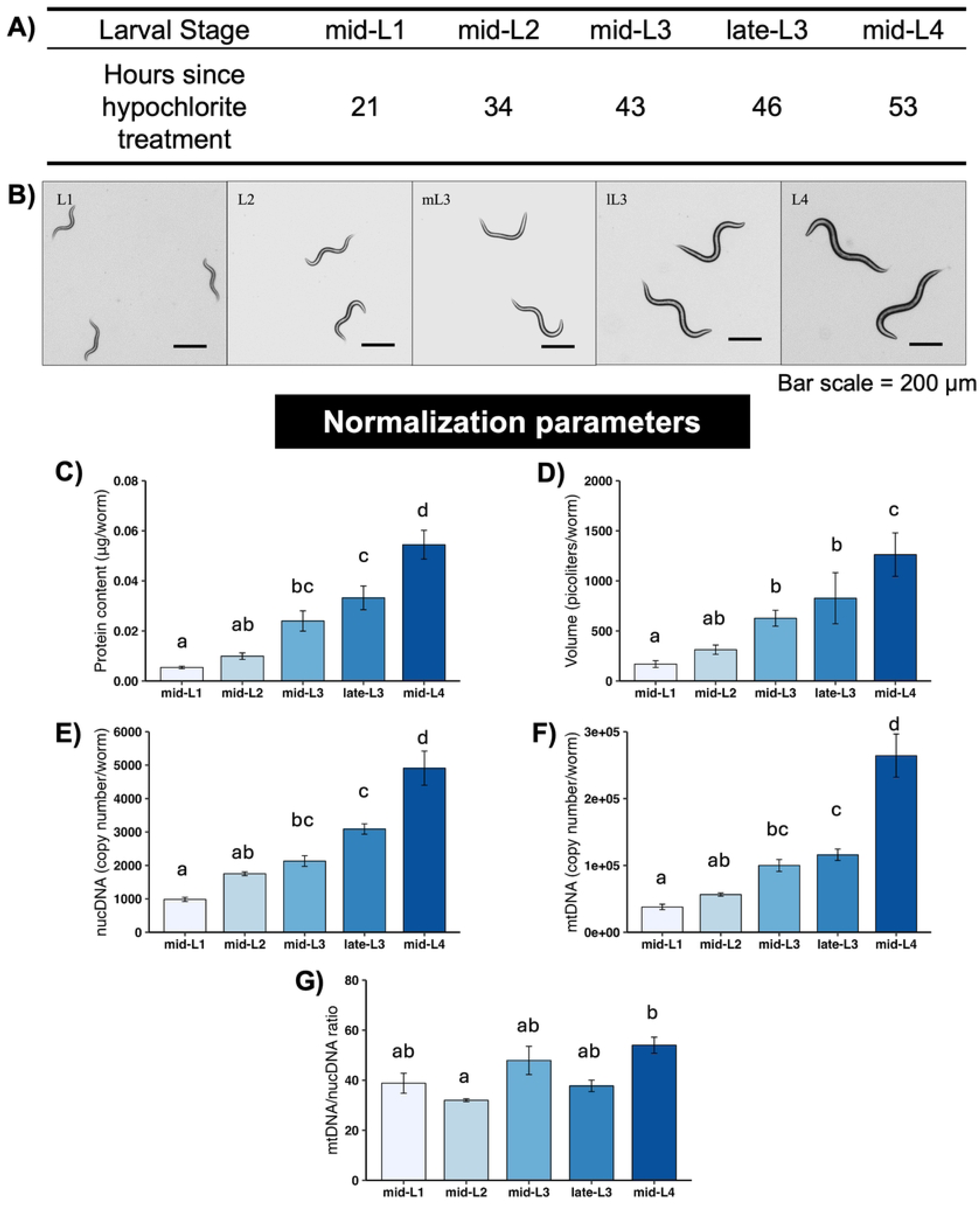
Normalization parameters for different larval stages of *C. elegans*. A) Larval stages and timepoints for each stage used in this study. B) Representative images of *C. elegans* at each stage. C) Total protein content at each stage. D) Worm volume at each stage. E) Nuclear DNA copy number at each stage. F) Mitochondrial DNA copy number at each stage. G) Mitochondrial/nuclear DNA ratio at each stage. All normalization endpoints had 3-5 biological replicates per stage. Different letters represent p < 0.05 with Tukey’s posthoc following ANOVA.

For each larval stage, for optimization, we tested final well concentrations of 8, 25 or 75 µM for FCCP, and 20, 40 or 80 µM for DCCD. We initially tested 20, 40, and 80 mM sodium azide, but found no additional inhibition of OCR and so used 10 mM used in all experiments shown here. We also tested 12.5, 50, and 90 µM FCCP in the L1 stage. Concentration optimization data for FCCP and DCCD are presented in **Figures S2-S3**. Using the criteria described above, we found that 25 µM FCCP was the best of the tested concentrations at all larval stages; 50 µM was also acceptable at L1 (but was not tested at other stages). 20 µM DCCD was consistently slow to achieve full inhibition of OCR, at all larval stages. 40 µM DCCD worked well at all larval stage except L4, where it was slow to take effect. 80 µM was best at L4, and may be acceptable at earlier larval stages, although in some cases 80 µM resulted in a puzzling waning of inhibition of OCR over time (**Fig. S3**). We note that while these concentrations were optimal among those we tested, under certain circumstances, by taking a subset of the top measurements after FCCP injection and lowest measurements after DCCD injection it may be possible to include data from a wider range of concentrations for comparisons between larval stages, increasing statistical power. For example, even in cases where FCCP or DCCD were slow to reach their full effect, by taking only the highest three and lowest two OCR values (respectively), it would be possible to measure FCCP-stimulated and DCCD-inhibited OCR accurately. Our most current, detailed protocol for measuring OCR using a Seahorse instrument is presented in **Supporting information file S4**. Note that full descriptions are provided for a 24-well instrument (used in this study), with some additional description relevant to using a 96-well instrument. Similarly, to maximize the utility of this protocol, we also include conditions for adult stages although we did not include adult data in this manuscript.

### 3.2 Use of parallel DCCD and FCCP injections

Because we previously found that FCCP injection altered the response to subsequent sodium azide injection at the L4 stage [37], we tested whether this would be true at other larval stages. Indeed, FCCP decreased the degree to which sodium azide would inhibit OCR at all larval stages except late L3 (**Fig. S5**). We also found smaller effects of DCCD and DMSO on the response to sodium azide injection (**Fig. S5**). Therefore, for the experiments in this manuscript, we injected DCCD and FCCP in parallel, using separate plates, rather than in sequence and on the same plate, as is more typically done for cell culture analyses. As a result, for each larval stage measurement, we had separate measurements for basal OCR (i.e., OCR measured prior to injection of either DCCD or FCCP). We tested statistically for whether there was a plate effect for the basal OCR measurements, and as expected, found none; therefore, we combined those results. However, we used plate-specific measurements for calculation of maximal OCR and spare capacity (FCCP plates) and ATP-linked and proton leak-associated OCR (DCCD plates). We calculated non-mitochondrial and mitochondrial basal OCR values using DCCD plates.

### 3.3 mtDNA count, but not mtDNA:ncDNA ratio, increases from mid-L1 to mid-L4 stages

The measured larval stage-specific parameters other than worm count that we used to normalize OCR results are shown in **Figure 1C-G**: total protein content, worm volume, mtDNA and ncDNA CN, and mtDNA:ncDNA ratio. Interestingly, mtDNA:ncDNA ratio only clearly increased at the L4 stage, and this increase was relatively modest (∼20%). An important aspect of *C. elegans* development that informs interpretation of these values for normalization purposes is that somatic cell divisions are invariant in all worms, and germ cell division limited through the L4 stage, such that ncDNA CN is essentially a readout for developmental progression.

### 3.4 Basal OCR

Basal OCR is simply the rate of total oxygen consumption measured in liquid without any additional injections or manipulations. It is generally comparable to the values obtained using older techniques such as Clark electrode-based measurements. Basal OCR increased consistently throughout development when results were normalized per worm or mtDNA/ncDNA ratio (**Fig. 2**). In contrast, when basal OCR is normalized to volume, total protein, ncDNA CN, or mtDNA CN, there is a 2- to 3-fold increase from L1 to L2, but little or no increase throughout the later developmental stages.

**Figure 2.**
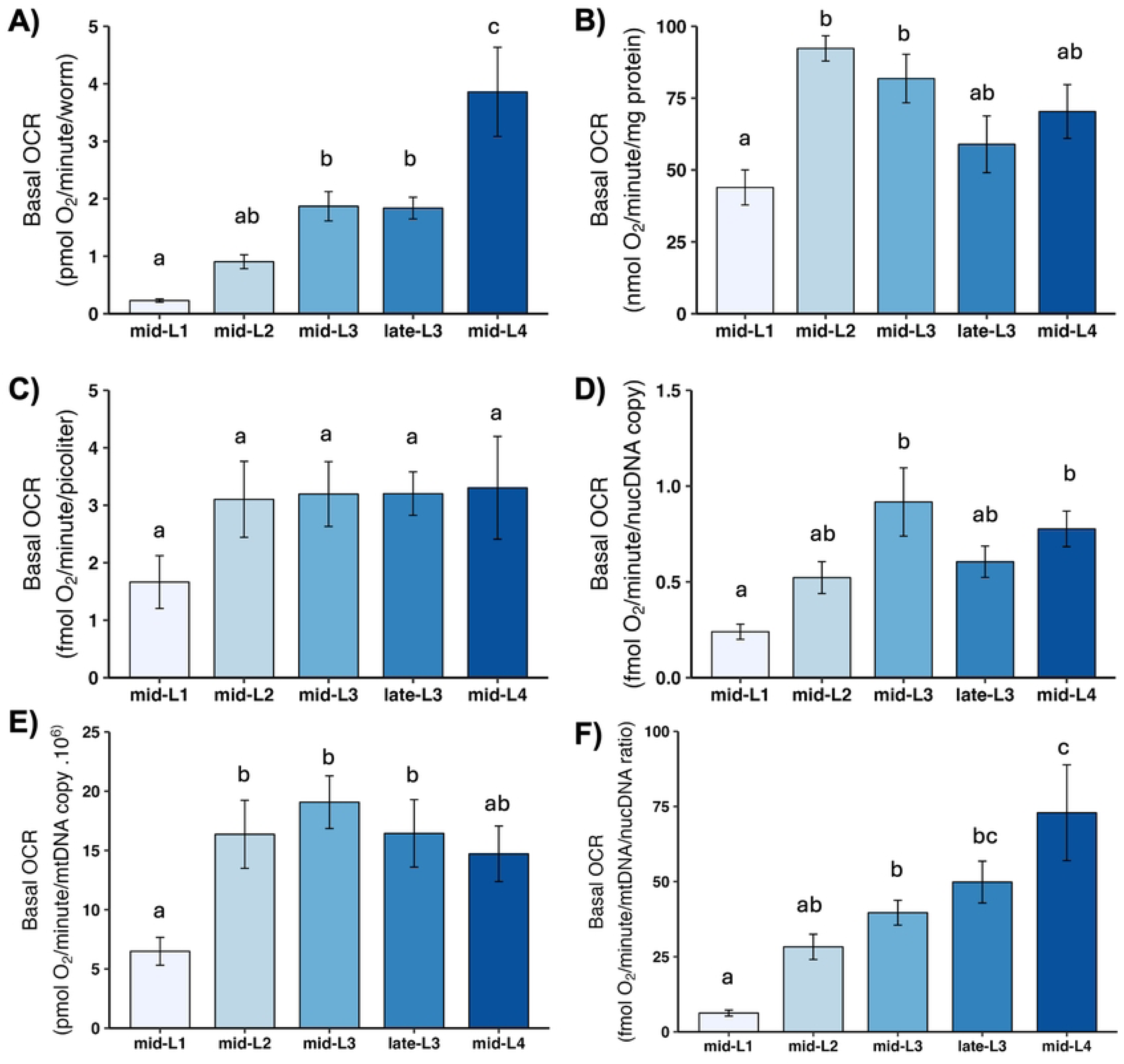
Basal oxygen consumption rate (OCR) at different larval stages after normalization. A) Basal OCR per worm. B) Basal OCR per mg protein. C) Basal OCR per volume. D) Basal OCR per ncDNA copy number. E) Basal OCR per mtDNA copy number. F) Basal OCR per mtDNA/ncDNA ratio. n = 3-5 biological replicates across stages. Different letters represent p < 0.05 with Tukey’s posthoc following ANOVA.

### 3.5 Basal mitochondrial OCR

Basal mitochondrial OCR is that portion of basal OCR that can be attributed to mitochondrial oxygen consumption (i.e., this excludes non-mitochondrial OCR). Although large variance resulted in a lack of statistically significant differences, the general pattern of mean mitochondrial OCR was essentially the same as observed for basal OCR: substantial increases across larval stages disappear, with the exception of an increase from L1 to L2, upon normalization to volume, total protein, ncDNA CN, or mtDNA CN (**Fig. 3**). This is highlighted by our calculation of the percent of total basal OCR that was mitochondrial: the percent was similar at all larval stages, 75-80% (**Fig. 3B**).

**Figure 3.**
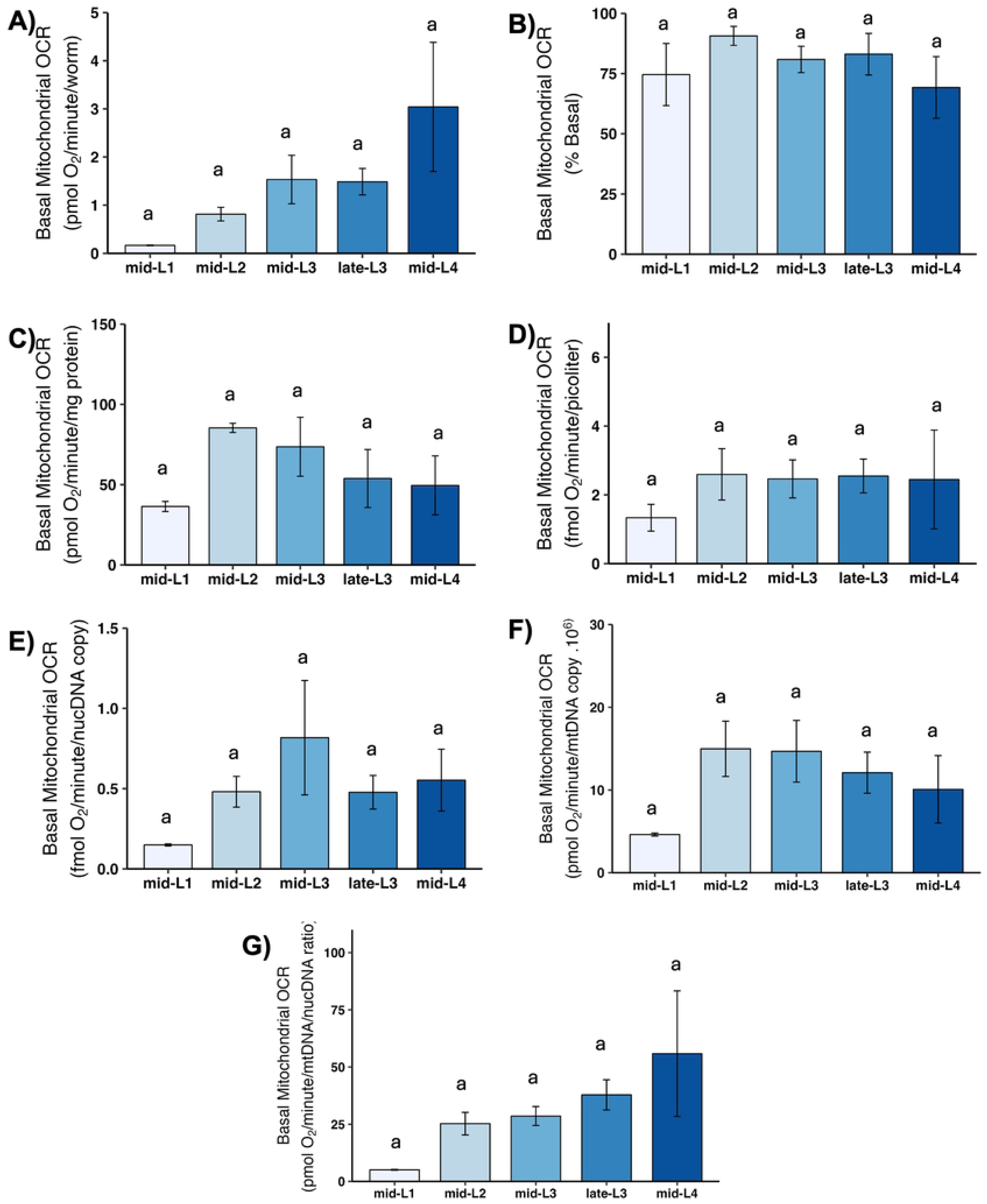
Basal mitochondrial oxygen consumption rate (OCR) at different larval stages after normalization. A) Basal mitochondrial OCR per worm. B) Basal mitochondrial OCR as percent total basal. C) Basal mitochondrial OCR per mg protein. D) Basal mitochondrial OCR per volume. E) Basal mitochondrial OCR per ncDNA copy number. F) Basal mitochondrial OCR per mtDNA copy number. G) Basal mitochondrial OCR per mtDNA/ncDNA ratio. n = 2-3 biological replicates across stages. Different letters represent p < 0.05 with Tukey’s posthoc following ANOVA.

### 3.6 ATP-linked OCR

ATP-linked OCR is defined experimentally as the amount of basal OCR that can be eliminated by inhibiting ATP synthase, and is intended to reflect the portion of mitochondrial oxygen consumption used to convert ADP to ATP—i.e., to “make energy.” Developmental changes in ATP-linked OCR were similar to those observed for total mitochondrial OCR (**Fig. 4**).

**Figure 4.**
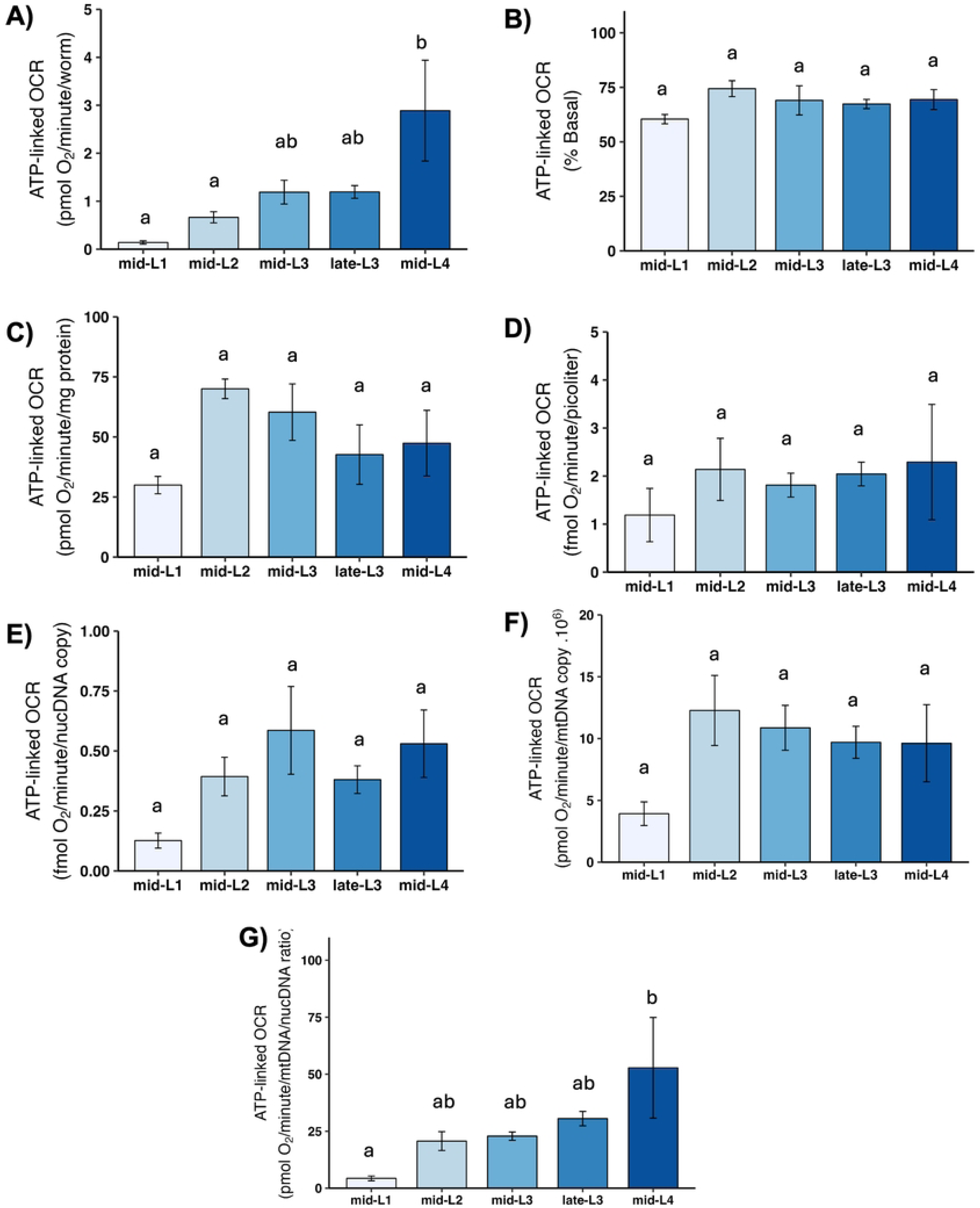
ATP-linked mitochondrial oxygen consumption rate (OCR) at different larval stages after normalization. A) ATP-linked OCR per worm. B) ATP-linked OCR as percent total basal. C) ATP-linked OCR per mg protein. D) ATP-linked OCR per volume. E) ATP-linked OCR per ncDNA copy number. F) ATP-linked OCR per mtDNA copy number. G) ATP-linked OCR per mtDNA/ncDNA ratio. Data shown using 40 µM DCCD for larval stages L1-late-L3 and 80 µM for L4 stage. n = 2-3 biological replicates across stages. Different letters represent p < 0.05 with Tukey’s posthoc following ANOVA.

### 3.7 Maximal OCR and spare respiratory capacity

Maximal OCR is determined by chemically uncoupling mitochondria; FCCP shuttles protons back into the matrix without generating ATP, which causes mitochondria to consume oxygen faster in order to maintain the proton gradient. Thus, maximal OCR serves as a measure of how quickly mitochondria can consume oxygen. The increase in OCR from the basal level is termed “spare respiratory capacity,” and reflects the ability of mitochondria to increase oxygen consumption on demand, which is important for stress response. We found a large increase in maximal OCR (**Fig. 5**) and spare respiratory capacity (**Fig. 6**) on per-worm basis, and a significant increase from L1 to L2 after other normalizations. A further increase occurred in most cases in one or both L3 stages, with a subsequent decrease in the L4 stage, although this was statistically significant only after normalization to mtDNA CN.

**Figure 5.**
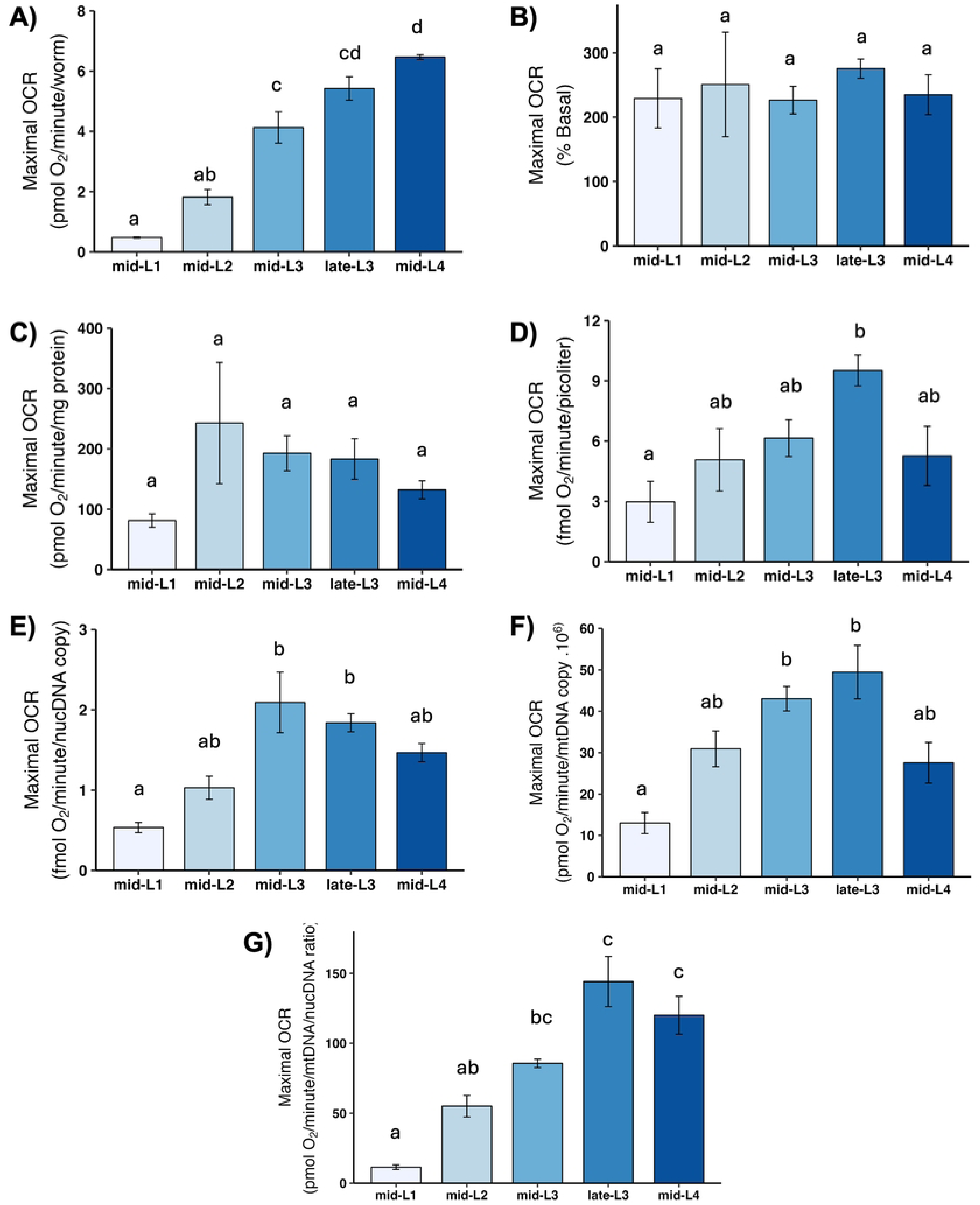
Maximal oxygen consumption rate (OCR) at different larval stages after normalization. A) Maximal OCR per worm. B) Maximal OCR as percent total basal. C) Maximal OCR per mg protein. D) Maximal OCR per volume. E) Maximal OCR per ncDNA copy number. F) Maximal OCR per mtDNA copy number. G) Maximal OCR per mtDNA/ncDNA ratio. Data shown using 25 µM FCCP across all stages. n = 2-3 biological replicates across stages. Different letters represent p < 0.05 with Tukey’s posthoc following ANOVA.

**Figure 6.**
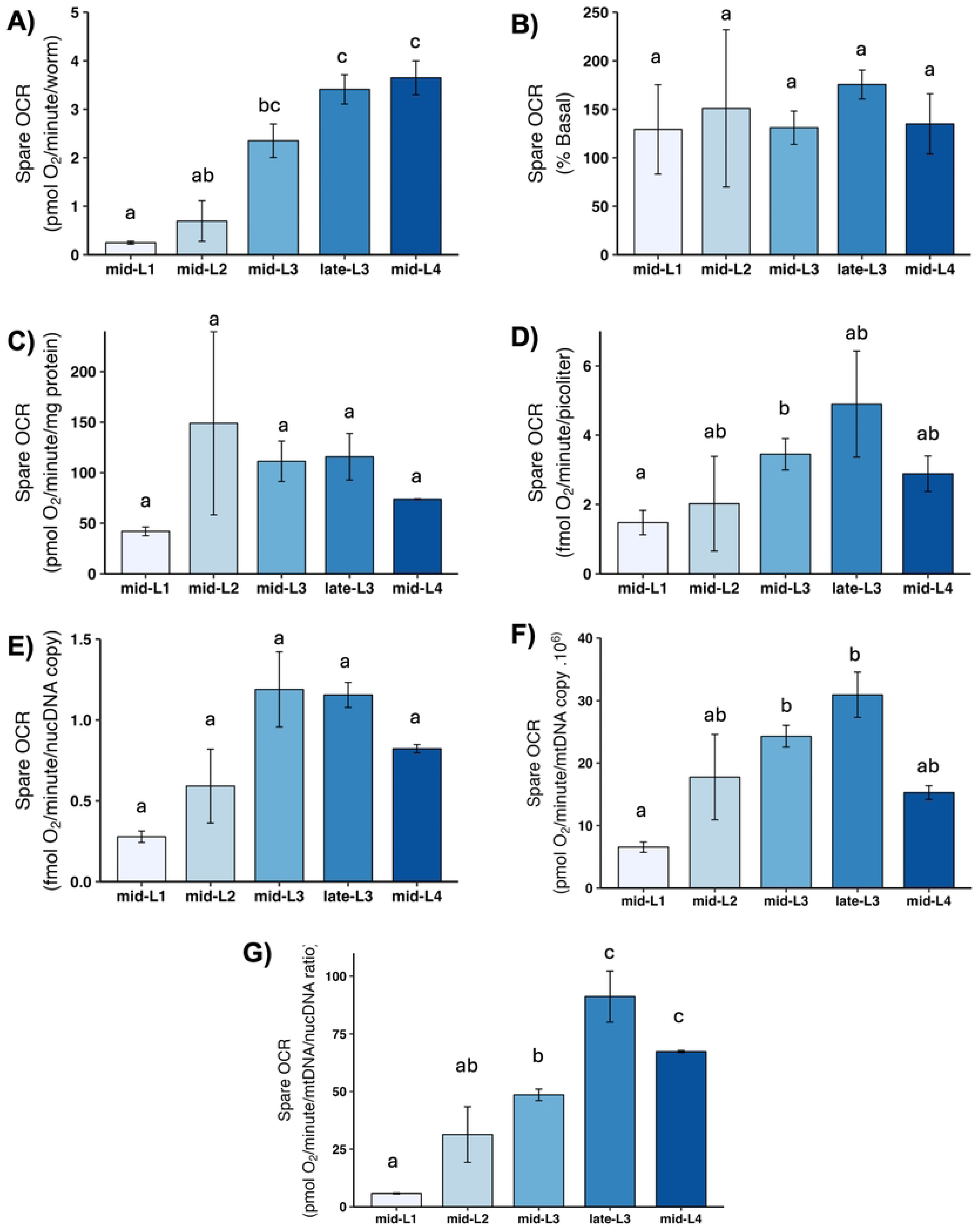
Spare respiratory capacity at different larval stages after normalization. A) Spare OCR per worm. B) Spare OCR as percent total basal. C) Spare OCR per mg protein. D) Spare OCR per volume. E) Spare OCR per ncDNA copy number. F) Spare OCR per mtDNA copy number. G) Spare OCR per mtDNA/ncDNA ratio. Data shown using 25 µM FCCP across all stages. n = 2-4 biological replicates across stages. Different letters represent p < 0.05 with Tukey’s posthoc following ANOVA.

### 3.8 Proton leak

“Proton leak” refers to oxygen consumed by mitochondria when ATP synthase is blocked. Because oxygen can only be consumed by mitochondria when there is electron flow, and electron flow is blocked if there is no dissipation of the proton gradient, any mitochondrial oxygen consumption that occurs when ATP synthase is blocked must reflect dissipation of the proton gradient by “leak.” “Leak” is thus defined as any such dissipation that is not mediated by ATP synthase, and includes important biological processes that may vary with developmental stage such as protein import, ion exchange, phosphate transport, transhydrogenase activity, and more [42]. We observed a pattern of apparent increase from L1 to L2, possibly peaking at L3 and then decreasing slightly upon normalization to factors other than worm count; however, these changes were not statistically significant (**Fig. 7**).

**Figure 7.**
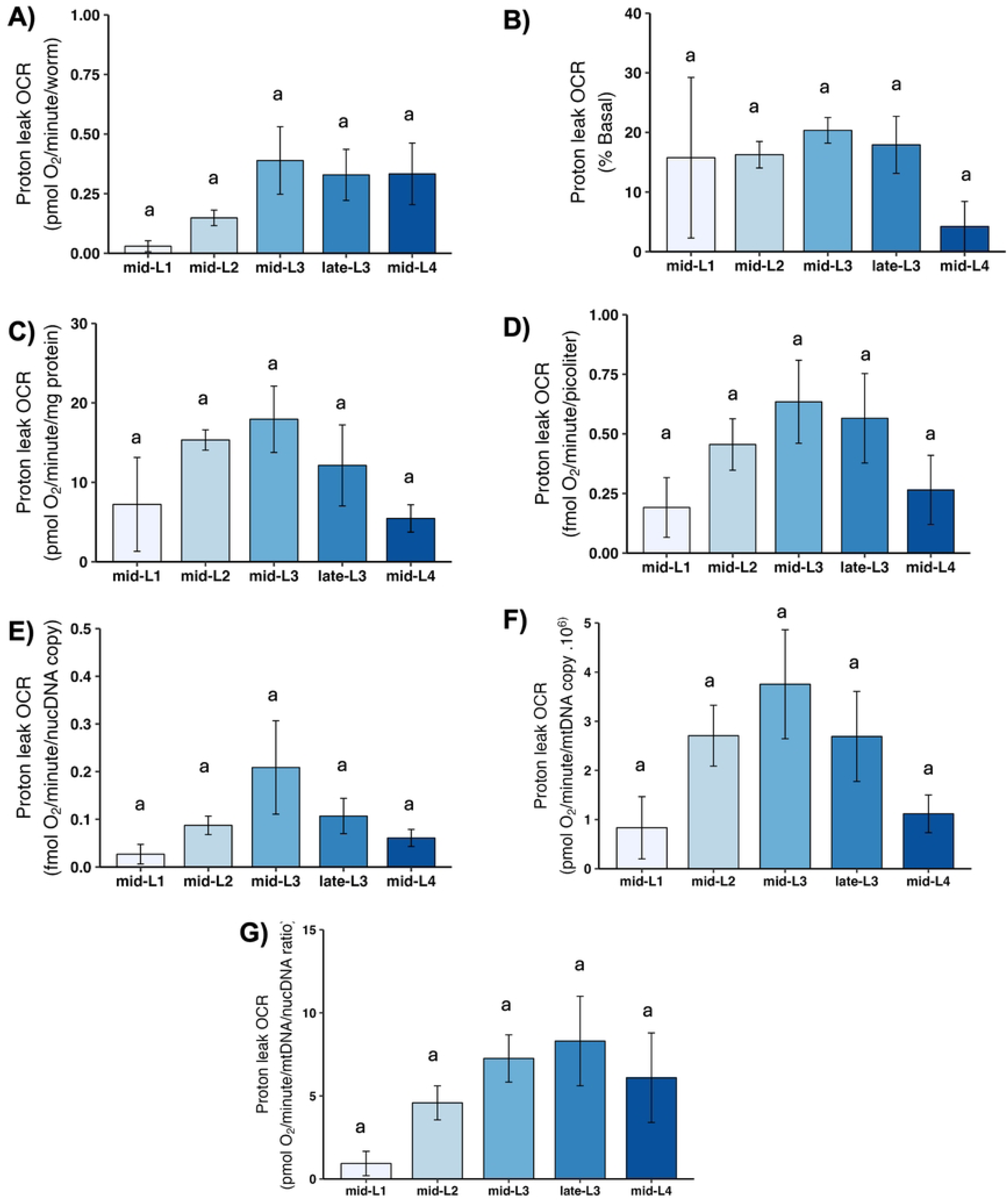
Proton leak at different larval stages after normalization. A) Proton leak OCR per worm. B) Proton leak OCR as percent total basal. C) Proton leak OCR per mg protein. D) Proton leak OCR per volume. E) Proton leak OCR per ncDNA copy number. F) Proton leak OCR per mtDNA copy number. G) Proton leak OCR per mtDNA/ncDNA ratio. n = 2-3 biological replicates across stages. Different letters represent p < 0.05 with Tukey’s posthoc following ANOVA.

### 3.9 ATP-linked OCR and Spare Respiratory Capacity as a function of total basal OCR

To test whether the proportion of total cellular OCR that is used by mitochondria changes as a function of developmental stage, we plotted these ratios (panels B in **Figures 3-7**). We observed no differences.

### 3.10 Non-mitochondrial OCR

Cellular processes other than the electron transport chain also consume oxygen; these include cytochrome P450 and other monooxygenases, NADPH oxidases, and other enzymes. Non-mitochondrial OCR is measured by inhibiting all oxidative phosphorylation using sodium azide, a potent complex IV inhibitor. Non-mitochondrial OCR increased substantially on a per-worm basis throughout development, but showed no change throughout development upon normalization by most other factors (**Fig. 8**). In particular, the pattern of an increase in mitochondrial OCR-related parameters from L1 to L2 was not observed for non-mitochondrial OCR.

**Figure 8.**
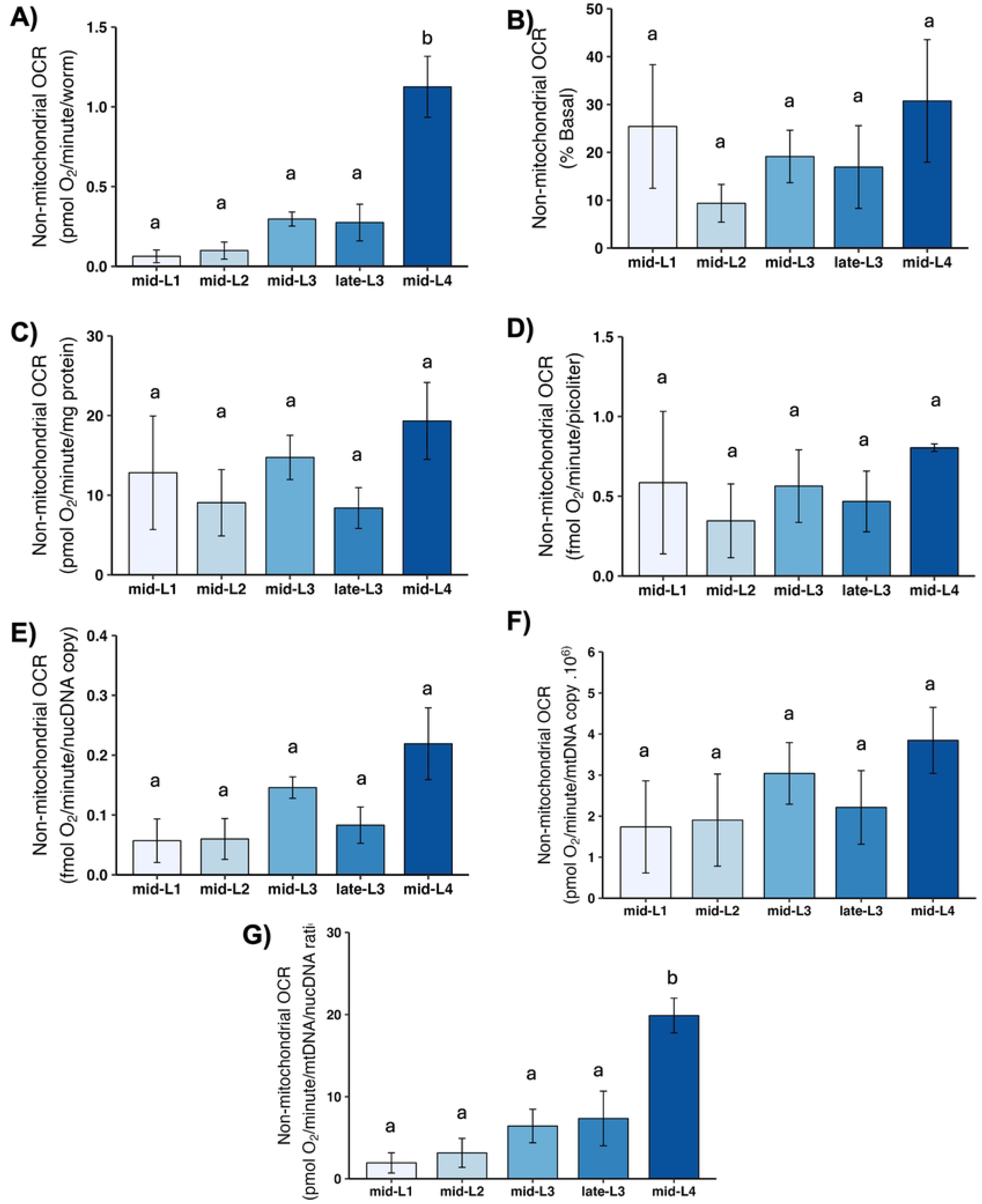
Non-mitochondrial oxygen consumption rate (OCR) at different larval stages after normalization. A) Non-mitochondrial OCR per worm. B) Non-mitochondrial OCR as percent total basal. C) Non-mitochondrial OCR per mg protein. D) Non-mitochondrial OCR per volume. E) Non-mitochondrial OCR per ncDNA copy number. F) Non-mitochondrial OCR per mtDNA copy number. G) Non-mitochondrial OCR per mtDNA/ncDNA ratio. n = 2-3 biological replicates across stages. Letters represent p < 0.05 with Tukey’s posthoc following ANOVA.

## 4. Discussion and conclusions

We report an optimized protocol for Seahorse-based analysis of OCR at all developmental stages in *C. elegans*. In the course of testing these parameters, we made two observations that are of biological interest. First, we learned that developmental trends in all measured OCR parameters vary qualitatively and quantitatively depending on how those values are normalized. Second, contrary to earlier ideas that most bioenergetic processes are anaerobic in early larval stages, we learned that there is significant oxidative phosphorylation occurring at all larval stages on a per-volume, per-protein, or per-mtDNA basis. Most of the normalized increase in oxidative phosphorylation that happens between the L1 and L4 stages occurs between L1 and L2, with a smaller increase observed in some parameters with some normalizations at the L3 stage. Basal OCR, maximal OCR, and spare respiratory capacity were slightly lower at the L4 than L3 stages upon normalization to mtDNA CN.

Very early developmental stages are reported to utilize energetic processes other than oxidative phosphorylation [43]. If and when this switch might occur in *C. elegans* has not been clearly defined. However, based on indirect evidence derived from a variety of phenotypes in which larval development is blocked or at least slowed when mtDNA replication, mitochondrial protein translation, or oxidative phosphorylation are inhibited, previous literature has suggested that the L3/L4 transition is accompanied by a metabolic switch from glycolysis to oxidative phosphorylation, or at least that inhibition of these processes signals for developmental delay or arrest at these (but not prior) stages [11–13, 29, 34]. This appeared consistent with a number of reports of dramatic increases in OCR through development when calculated on a per-nematode basis, as our results also show. Furthermore, Vanfleteren and De Vreese [32] reported peak protein-normalized OCR at the L3 and L4 stages, again consistent with a required metabolic switch at this stage. On the other hand, Houthoofd et al. reported peak basal respiration at the L2 stage, after normalization to protein content [33]. Our results clearly show significant oxidative phosphorylation at L1, a strong increase at L2, and possibly some additional increase at L3, depending on how normalization is done. These results suggest that however worms sense mitochondrial dysfunction and therefore arrest at L3, it is unlikely to be failure to activate oxidative phosphorylation, as any such inhibition should be able to be sensed by L2 or even L1 stage worms. One caveat to this conclusion is that our measurements were made at the whole-organism level and therefore cannot exclude a cell-specific oxidative phosphorylation-sensing mechanism. A second caveat is that it is possible that constraint of maximal (but not basal mitochondrial or ATP-linked) OCR could somehow be limiting for transition to L4. Such a constraint could be effective at the L3 to L4 stage because the maximal OCR is somewhat reduced in L4 compared to L3, in particular when normalized to mtDNA, which we assume serves as a proxy for amount of mitochondria. Another biological aspect of the L3 to L4 transition that could relate to developmental arrest is the onset of gonadal development and gametogenesis [44]. Cellular reactive oxygen species (ROS) and redox status are essential for proliferation, differentiation, and migration, and mitochondrial ROS production has a major role in the control of such processes [45]. It is possible that the L3 arrest upon mitochondrial dysfunction may be due to a strict control of the redox status necessary for proper gametogenesis. Worms have a more reducing environment during the L3 to L4 transition [46] and dysfunctional mitochondrial may produce higher levels of ROS [47]. Thus, in the light of such findings, our results suggests that the L3 arrest upon mitochondrial dysfunction could result from signaling events or the necessity of a strict redox status control rather than a reliance of oxidative phosphorylation for energy production starting only at this developmental stage. We recommend future studies focusing on a deeper understanding of the mitochondrial control of *C. elegans* cellular redox status and development.

Choice of normalization parameters is critical. One approach is to normalize OCR to volume rather than number of worms. This is important even when analyzing OCR in cell culture [48], where cell size varies with cell cycle but to a much smaller degree than is the case for developing worms. From just-hatched L1s to late-L4 worms, there is nearly a 10-fold increase in volume [49](our own results show a slightly smaller change, likely because we examined a slightly reduced developmental time course). Thus, worm number is not a useful normalization value unless all worms in the experiment are nearly identical in size. Note that even within larval stages there is significant growth [49], as is also true in adulthood. A second normalization approach is to protein content, either total or mitochondrial. This is helpful, although more work-intensive. A third option is to normalize to ncDNA CN, which serves as a proxy for cell number and developmental stage at least during development in *C. elegans*, since somatic cell development is identical in every individual in this species. This approach also fails to take into account cell size variation, however. It also works less well in gravid adults, where a high proportion of nuclear genomes comes from the germline, such that effects of experimental manipulations on OCR could be the indirect result of how those manipulations affect germ cell division rather than how they affect mitochondria function *per se*. Normalizing OCR to volume or ncDNA CN has the disadvantage of not demonstrating whether a change in OCR is a function of more or fewer mitochondria per worm or per cell, versus a function of the same number of mitochondria not consuming the same amount of oxygen as controls (e.g., because of a genetic deficiency or chemical effect). For this purpose, we recommend measuring mtDNA CN, although it is also possible (but more challenging) to measure total mitochondrial protein, or specific mitochondrial proteins, or other proxies for mitochondrial “amount.” Such an approach allowed us to show that in worms, exercise improved mitochondrial function as measured by OCR and other parameters, without altering mtDNA CN, suggesting improved mitochondrial function on a per-mitochondrion basis [26]. Ultimately, which of these is chosen will depend on the experimental questions being asked; often, it will be valuable to compare the result of multiple normalization procedures. Of note, these considerations are also important in circumstances outside those examined in this study. For example, mtDNA CN is also regulated by age in adults [50] and starvation [41], and so changes in OCR in those contexts need to be interpreted in the context of changes in mitochondrial content.

In conclusion, we present a carefully optimized protocol for Seahorse-based analysis of oxygen consumption rate (OCR) throughout the developmental stages of *C. elegans*, and highlight the importance of careful parameter selection for normalization in such analyses. Moreover, our study enhances our understanding of *C. elegans* physiology throughout development and provides valuable insights applicable to broader contexts, including significant MRC-mediated oxygen consumption in all larval stages, especially after L1. Future work measuring gamete and embryonic metabolism, changes with age, and cell type-specific metabolism will further increase our understanding of how mitochondrial metabolism contributes to worm development, physiological functions, stress response, and aging.

## Author contributions

Conceptualization: DFM, CMB, JNM; Formal analysis: DFM, CMB, JNM; Funding acquisition, resources, and supervision: JNM; Investigation: DFM, LP, ITR; Methodology: DFM, CMB, KSM, JNM; Visualization: DFM, CMB; Writing: LP, DFM, KSM, JNM.

## Acknowledgments

This work was funded by the National Institute of Health (R01ES028218, P42ES010356, R01ES034270). Some strains were provided by the Caenorhabditis Genetics Center, which is funded by NIH Office of Research Infrastructure Programs (P40 OD010440).

## Supporting information

**Excel file containing all Seahorse, DNA copy number, volume, and protein analysis data.**

**S2.**
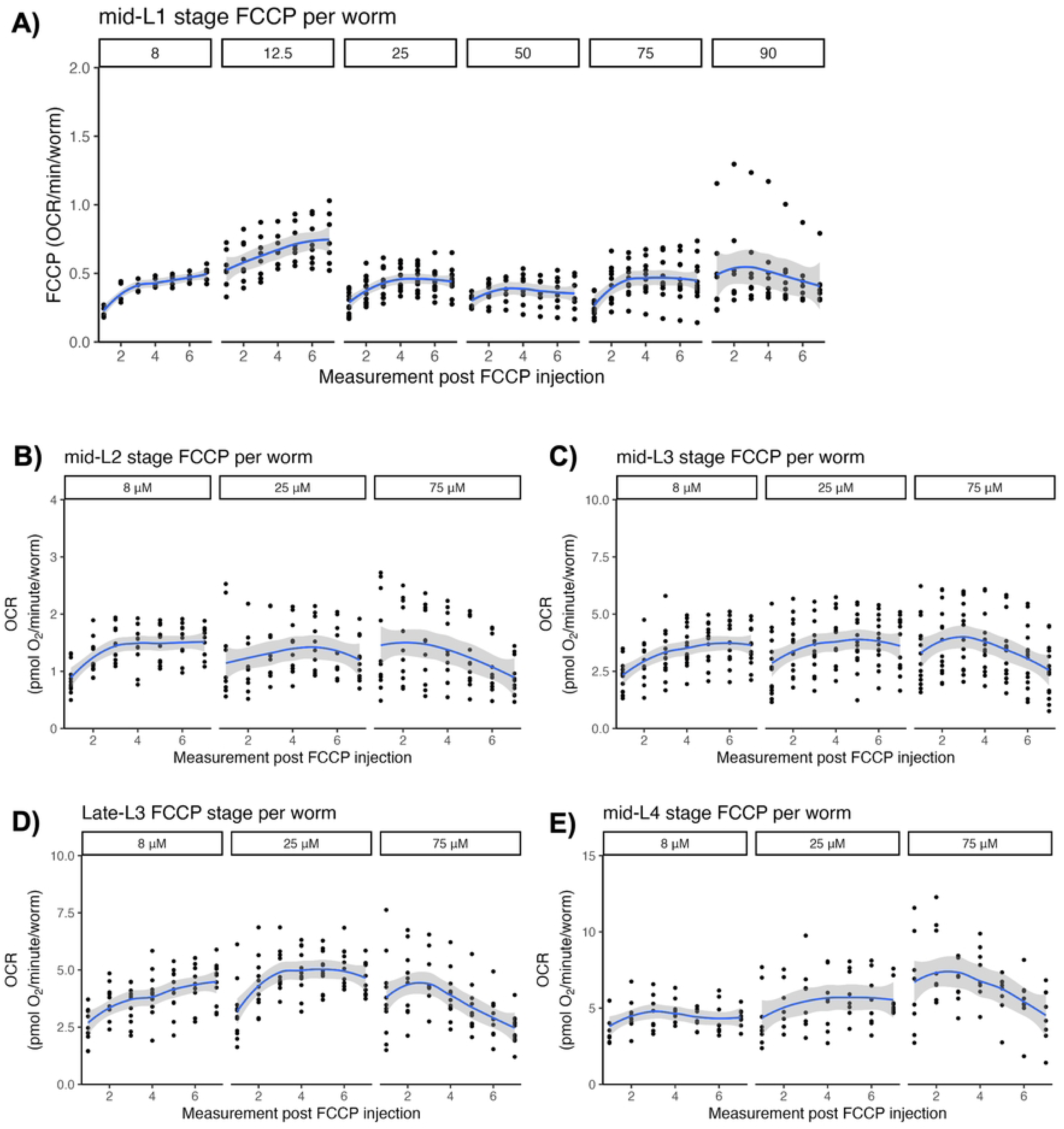
Figure S2 shows oxygen consumption rates at the L1 (panel A), L2 (panel B), mid-L3 (panel C), late L3 (panel D), and L4 (panel E) stages at different timepoints after the injection of different concentrations of FCCP, normalized per worm. Blue line represents local polynomial regression fitting of the data for visualization with 95% confidence interval using the geom_smooth function in ggplot2 package in R version 4.2.1. Figures A-E represent data across 1-4 biological replicates with 3-6 technical replicates (L1 – 1-3 biological reps with 4-5 technical replicates; L2 – three biological reps with 3-5 technical replicates; L3 – four biological reps with 3-5 technical replicates; Late-L3-three biological reps 3-5 technical replicates; L4 – two biological reps with 3-5 technical replicates).

**S3.**
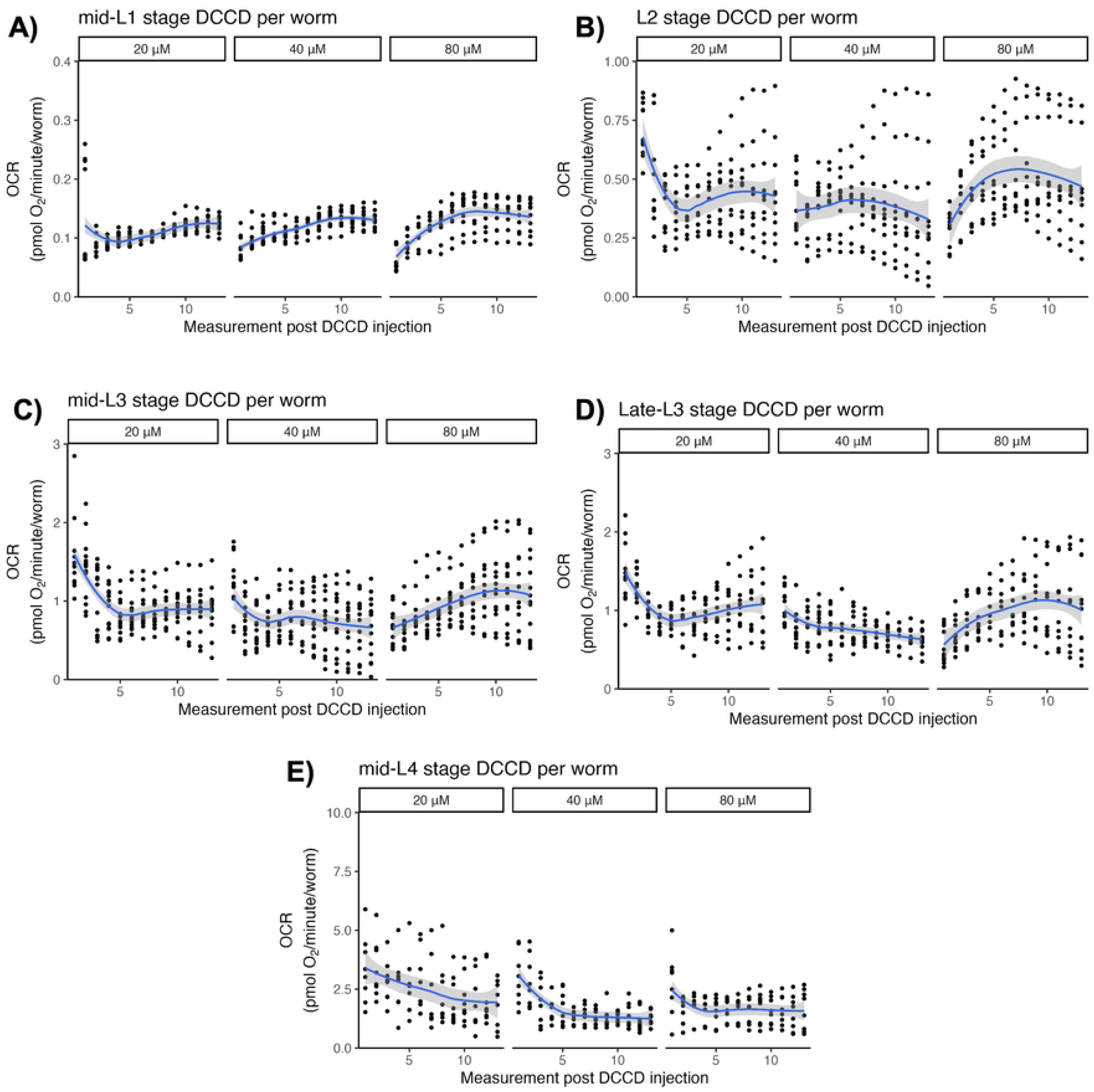
Figure S3 shows oxygen consumption rates at the L1 (panel A), L2 (panel B), mid-L3 (panel C), late L3 (panel D), and L4 (panel E) stages at different timepoints after the injection of different concentrations of DCCD, normalized per worm. Blue line represents local polynomial regression fitting of the data for visualization with 95% confidence interval using the geom_smooth function in ggplot2 package in R version 4.2.1. Figures A-E represent data across 2-4 biological replicates with 3-6 technical replicates (L1 – two biological reps with 4-6 technical reps; L2 – three biological reps with 3-5 technical reps; L3 – four biological reps with 3-5 technical reps; Late-L3-three biological reps with 3-5 technical reps; L4 – two biological reps with 3-5 technical reps).

**S4. Word document with our laboratory Seahorse protocol.** Please note that this protocol includes information for adult lifestages as well and larval lifestages, and information about using a 96-instead of 24-well Seahorse instrument.

**S5.**
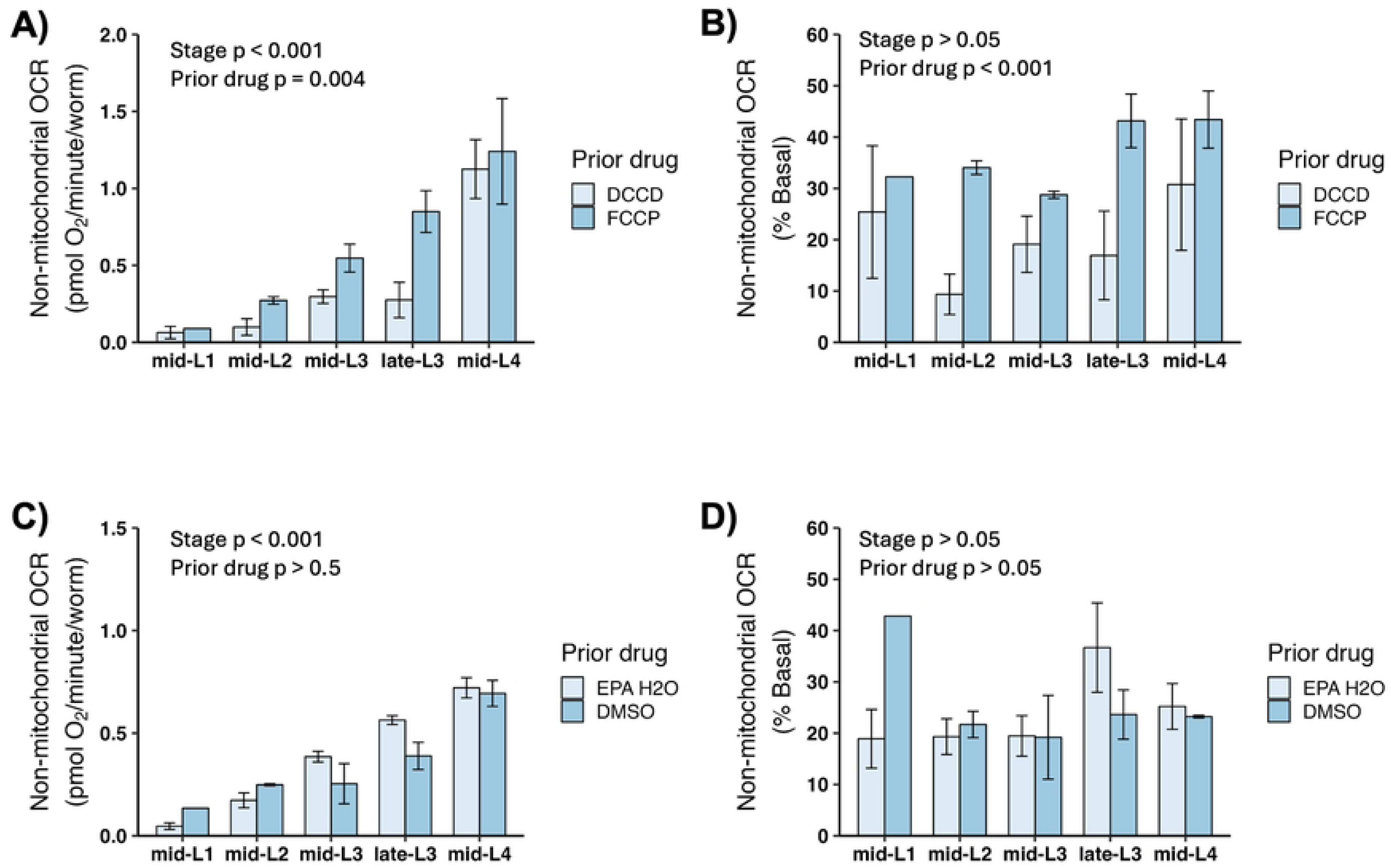
Figure S5 shows the effects of including DMSO solvent vs EPA water, prior FCCP injections, and prior DCCD injections on sodium azide-mediated inhibition of oxygen consumption rates at different larval stages. Non-mitochondrial OCR normalized per worm and percent basal after injection with FCCP and DCCD. n = 1-4 biological replicates, p-values from two-way ANOVA.

## References

1. Naviaux RK. Perspective: Cell danger response Biology-The new science that connects environmental health with mitochondria and the rising tide of chronic illness. Mitochondrion. 2020;51:40–5. Epub 20191223. doi: 10.1016/j.mito.2019.12.005. PubMed PMID: 31877376.

2. Nunnari J, Suomalainen A. Mitochondria: in sickness and in health. Cell. 2012;148(6):1145–59. Epub 2012/03/20. doi: 10.1016/j.cell.2012.02.035. PubMed PMID: 22424226.

3. Suomalainen A, Battersby BJ. Mitochondrial diseases: the contribution of organelle stress responses to pathology. Nat Rev Mol Cell Biol. 2018;19(2):77–92. Epub 20170809. doi: 10.1038/nrm.2017.66. PubMed PMID: 28792006.

4. Lightowlers RN, Taylor RW, Turnbull DM. Mutations causing mitochondrial disease: What is new and what challenges remain? Science. 2015;349(6255):1494–9. Epub 2015/09/26. doi: 10.1126/science.aac7516. PubMed PMID: 26404827.

5. Meyer JN, Leung MC, Rooney JP, Sendoel A, Hengartner MO, Kisby GE, et al. Mitochondria as a target of environmental toxicants. Toxicol Sci. 2013;134(1):1–17. Epub 2013/05/01. doi: 10.1093/toxsci/kft102. PubMed PMID: 23629515; PubMed Central PMCID: PMCPMC3693132.

6. Cohen BH. Pharmacologic effects on mitochondrial function. Developmental disabilities research reviews. 2010;16(2):189–99. Epub 2010/09/08. doi: 10.1002/ddrr.106. PubMed PMID: 20818734.

7. Picard M, Shirihai OS. Mitochondrial signal transduction. Cell Metab. 2022;34(11):1620–53. doi: 10.1016/j.cmet.2022.10.008. PubMed PMID: 36323233; PubMed Central PMCID: PMCPMC9692202.

8. Corsi AK, Wightman B, Chalfie M. A Transparent Window into Biology: A Primer on Caenorhabditis elegans. Genetics. 2015;200(2):387–407. doi: 10.1534/genetics.115.176099. PubMed PMID: WOS:000356509100001.

9. Maglioni S, Ventura N. C. elegans as a model organism for human mitochondrial associated disorders. Mitochondrion. 2016. Epub 2016/02/26. doi: 10.1016/j.mito.2016.02.003. PubMed PMID: 26906059.

10. Maurer LL, Luz AL, Meyer JN. Detection of mitochondrial toxicity of environmental pollutants using Caenorhabditis elegans. In: Dykens JA, Will Y, editors. Drug-Induced Mitochondrial Dysfunction: Wiley; 2018.

11. Tsang WY, Sayles LC, Grad LI, Pilgrim DB, Lemire BD. Mitochondrial respiratory chain deficiency in Caenorhabditis elegans results in developmental arrest and increased life span. The Journal of biological chemistry. 2001;276(34):32240–6. PubMed PMID: 11410594.

12. Tsang WY, Lemire BD. Mitochondrial genome content is regulated during nematode development. Biochem Biophys Res Commun. 2002;291(1):8–16. PubMed PMID: 11829454.

13. Bratic I, Hench J, Henriksson J, Antebi A, Burglin TR, Trifunovic A. Mitochondrial DNA level, but not active replicase, is essential for Caenorhabditis elegans development. Nucleic acids research. 2009;37(6):1817–28. PubMed PMID: 19181702.

14. Bess AS, Crocker TL, Ryde IT, Meyer JN. Mitochondrial dynamics and autophagy aid in removal of persistent mitochondrial DNA damage in Caenorhabditis elegans. Nucleic acids research. 2012. Epub 2012/06/22. doi: 10.1093/nar/gks532. PubMed PMID: 22718972.

15. Luz AL, Godebo TR, Smith LL, Leuthner TC, Maurer LL, Meyer JN. Deficiencies in mitochondrial dynamics sensitize Caenorhabditis elegans to arsenite and other mitochondrial toxicants by reducing mitochondrial adaptability. Toxicology. 2017. doi: 10.1016/j.tox.2017.05.018. PubMed PMID: 28602540.

16. Maglioni S, Mello DF, Schiavi A, Meyer JN, Ventura N. Mitochondrial bioenergetic changes during development as an indicator of C. elegans health-span. Aging (Albany NY). 2019;11(16):6535–54. Epub 2019/08/28. doi: 10.18632/aging.102208. PubMed PMID: 31454791; PubMed Central PMCID: PMCPMC6738431.

17. Rea SL, Ventura N, Johnson TE. Relationship between mitochondrial electron transport chain dysfunction, development, and life extension in Caenorhabditis elegans. PLoS Biol. 2007;5(10):e259. PubMed PMID: 17914900.

18. Dillin A, Hsu AL, Arantes-Oliveira N, Lehrer-Graiwer J, Hsin H, Fraser AG, et al. Rates of behavior and aging specified by mitochondrial function during development. Science. 2002;298(5602):2398-401. PubMed PMID: 12471266.

19. Hershberger KA, Rooney JP, Turner EA, Donoghue LJ, Bodhicharla R, Maurer LL, et al. Early-life mitochondrial DNA damage results in lifelong deficits in energy production mediated by redox signaling in Caenorhabditis elegans. Redox Biol. 2021;43:102000. Epub 2021/05/17. doi: 10.1016/j.redox.2021.102000. PubMed PMID: 33993056.

20. Cypser JR, Johnson TE. Multiple stressors in *Caenorhabditis elegans* induce stress hormesis and extended longevity. Journals of Gerontology Series A-Biological Sciences and Medical Sciences. 2002;57(3):B109–B14. PubMed PMID: ISI:000174206900004.

21. Caito SW, Aschner M. NAD+ Supplementation Attenuates Methylmercury Dopaminergic and Mitochondrial Toxicity in Caenorhabditis Elegans. Toxicol Sci. 2016;151(1):139–49. Epub 20160210. doi: 10.1093/toxsci/kfw030. PubMed PMID: 26865665; PubMed Central PMCID: PMCPMC4914800.

22. Gonzalez-Hunt CP, Leung MC, Bodhicharla RK, McKeever MG, Arrant AE, Margillo KM, et al. Exposure to mitochondrial genotoxins and dopaminergic neurodegeneration in Caenorhabditis elegans. PloS one. 2014;9(12):e114459. Epub 20141208. doi: 10.1371/journal.pone.0114459. PubMed PMID: 25486066; PubMed Central PMCID: PMCPMC4259338.

23. Mao K, Breen P, Ruvkun G. Mitochondrial dysfunction induces RNA interference in C. elegans through a pathway homologous to the mammalian RIG-I antiviral response. PLoS Biol. 2020;18(12):e3000996. Epub 20201202. doi: 10.1371/journal.pbio.3000996. PubMed PMID: 33264285; PubMed Central PMCID: PMCPMC7735679.

24. Mello DF, Bergemann CM, Fisher K, Chitrakar R, Bijwadia SR, Wang Y, et al. Rotenone Modulates Caenorhabditis elegans Immunometabolism and Pathogen Susceptibility. Front Immunol. 2022;13:840272. Epub 20220222. doi: 10.3389/fimmu.2022.840272. PubMed PMID: 35273616; PubMed Central PMCID: PMCPMC8902048.

25. Guha S, Mathew ND, Konkwo C, Ostrovsky J, Kwon YJ, Polyak E, et al. Combinatorial glucose, nicotinic acid and N-acetylcysteine therapy has synergistic effect in preclinical C. elegans and zebrafish models of mitochondrial complex I disease. Hum Mol Genet. 2021;30(7):536–51. doi: 10.1093/hmg/ddab059. PubMed PMID: 33640978; PubMed Central PMCID: PMCPMC8120136.

26. Hartman JH, Smith LL, Gordon KL, Laranjeiro R, Driscoll M, Sherwood DR, et al. Swimming Exercise and Transient Food Deprivation in Caenorhabditis elegans Promote Mitochondrial Maintenance and Protect Against Chemical-Induced Mitotoxicity. Sci Rep. 2018;8(1):8359. Epub 20180529. doi: 10.1038/s41598-018-26552-9. PubMed PMID: 29844465; PubMed Central PMCID: PMCPMC5974391.

27. Luz AL, Godebo TR, Bhatt DP, Ilkayeva OR, Maurer LL, Hirschey MD, et al. From the Cover: Arsenite Uncouples Mitochondrial Respiration and Induces a Warburg-like Effect in Caenorhabditis elegans. Toxicol Sci. 2016;152(2):349–62. Epub 20160520. doi: 10.1093/toxsci/kfw093. PubMed PMID: 27208080; PubMed Central PMCID: PMCPMC4960910.

28. Zhang H, Burr SP, Chinnery PF. The mitochondrial DNA genetic bottleneck: inheritance and beyond. Essays Biochem. 2018;62(3):225–34. Epub 20180720. doi: 10.1042/EBC20170096. PubMed PMID: 29880721.

29. Leung MC, Rooney JP, Ryde IT, Bernal AJ, Bess AS, Crocker TL, et al. Effects of early life exposure to ultraviolet C radiation on mitochondrial DNA content, transcription, ATP production, and oxygen consumption in developing Caenorhabditis elegans. BMC pharmacology & toxicology. 2013;14:9. Epub 2013/02/05. doi: 10.1186/2050-6511-14-9. PubMed PMID: 23374645; PubMed Central PMCID: PMC3621653.

30. Bess AS, Leung MC, Ryde IT, Rooney JP, Hinton DE, Meyer JN. Effects of mutations in mitochondrial dynamics-related genes on the mitochondrial response to ultraviolet C radiation in developing Caenorhabditis elegans. Worm. 2013;2(1):e23763. doi: 10.4161/worm.23763. PubMed PMID: 24058863; PubMed Central PMCID: PMCPMC3670464.

31. Harvey AJ. Mitochondria in early development: linking the microenvironment, metabolism and the epigenome. Reproduction. 2019;157(5):R159–R79. doi: 10.1530/REP-18-0431. PubMed PMID: 30870807.

32. Vanfleteren JR, De Vreese A. Rate of aerobic metabolism and superoxide production rate potential in the nematode Caenorhabditis elegans. J Exp Zool. 1996;274(2):93–100. doi: 10.1002/(SICI)1097-010X(19960201)274:2<93::AID-JEZ2>3.0.CO;2-8. PubMed PMID: 8742689.

33. Houthoofd K, Braeckman BP, Lenaerts I, Brys K, De Vreese A, Van Eygen S, et al. Ageing is reversed, and metabolism is reset to young levels in recovering dauer larvae of C. elegans. Exp Gerontol. 2002;37(8-9):1015–21. PubMed PMID: 12213552.

34. Huang SH, Lin YW. Bioenergetic Health Assessment of a Single Caenorhabditis elegans from Postembryonic Development to Aging Stages via Monitoring Changes in the Oxygen Consumption Rate within a Microfluidic Device. Sensors (Basel). 2018;18(8). Epub 20180728. doi: 10.3390/s18082453. PubMed PMID: 30060586; PubMed Central PMCID: PMCPMC6111518.

35. Lewis JA, Fleming JT. Basic Culture Methods. In: Epstein HF, Shakes DC, editors. *Caenorhabditis elegans:* Modern Biological Analysis of an Organism. San Digo, CA: Academic Press; 1995. p. 3–29.

36. Williams PL, Dusenbery DB. Using the nematode Caenorhabditis elegans to predict mammalian acute lethality to metallic salts. Toxicol Ind Health. 1988;4(4):469–78. doi: 10.1177/074823378800400406. PubMed PMID: 3188044.

37. Luz AL, Rooney JP, Kubik LL, Gonzalez CP, Song DH, Meyer JN. Mitochondrial Morphology and Fundamental Parameters of the Mitochondrial Respiratory Chain Are Altered in Caenorhabditis elegans Strains Deficient in Mitochondrial Dynamics and Homeostasis Processes. PloS one. 2015;10(6):e0130940. Epub 2015/06/25. doi: 10.1371/journal.pone.0130940. PubMed PMID: 26106885; PubMed Central PMCID: PMC4480853.

38. Luz AL, Smith LL, Rooney JP, Meyer JN. Seahorse Extracellular Flux-based analysis of cellular respiration in Caenorhabditis elegans. Current Protocols in Toxicology. 2015;66:25.7.1-.7.15.

39. Rooney JP, Ryde IT, Saunders LH, Colton MD, Germ KE, Mayer GD, et al. PCR-based determination of mitochondrial DNA copy number in multiple species. Methods in Molecular Biology: Mitochondrial Regulation: Methods and Protocols. 2015.

40. Moore BT, Jordan JM, Baugh LR. WormSizer: high-throughput analysis of nematode size and shape. PloS one. 2013;8(2):e57142. Epub 20130222. doi: 10.1371/journal.pone.0057142. PubMed PMID: 23451165; PubMed Central PMCID: PMCPMC3579787.

41. Hibshman JD, Leuthner TC, Shoben C, Mello DF, Sherwood DR, Meyer JN, et al. Nonselective autophagy reduces mitochondrial content during starvation in Caenorhabditis elegans. Am J Physiol Cell Physiol. 2018;315(6):C781–C92. Epub 2018/08/23. doi: 10.1152/ajpcell.00109.2018. PubMed PMID: 30133321; PubMed Central PMCID: PMCPMC6336938.

42. Nicholls DG, Ferguson SJ. The chemiosmotic proton circuit in isolated organelles. Bioenergetics 4. New York: Academic Press; 2013. p. 53–88.

43. Knudsen TB, Green ML. Response characteristics of the mitochondrial DNA genome in developmental health and disease. Birth Defects Res C Embryo Today. 2004;72(4):313–29. doi: 10.1002/bdrc.20028. PubMed PMID: 15662705.

44. Pazdernik N, Schedl T. Introduction to germ cell development in Caenorhabditis elegans. Adv Exp Med Biol. 2013;757:1–16. doi: 10.1007/978-1-4614-4015-4_1. PubMed PMID: 22872472; PubMed Central PMCID: PMCPMC3781019.

45. Sena LA, Chandel NS. Physiological roles of mitochondrial reactive oxygen species. Mol Cell. 2012;48(2):158–67. doi: 10.1016/j.molcel.2012.09.025. PubMed PMID: 23102266; PubMed Central PMCID: PMCPMC3484374.

46. Back P, De Vos WH, Depuydt GG, Matthijssens F, Vanfleteren JR, Braeckman BP. Exploring real-time in vivo redox biology of developing and aging Caenorhabditis elegans. Free Radic Biol Med. 2012;52(5):850–9. Epub 20111223. doi: 10.1016/j.freeradbiomed.2011.11.037. PubMed PMID: 22226831.

47. Timme-Laragy AR, Di Giulio RT, Meyer JN. Reactive Oxygen Species and Redox Stress. In: Willett KL, Aluru N, editors. Toxicology of Fishes. 2 ed. New York: CRC Press; 2024. p. 121–45.

48. Renner K, Amberger A, Konwalinka G, Kofler R, Gnaiger E. Changes of mitochondrial respiration, mitochondrial content and cell size after induction of apoptosis in leukemia cells. Biochim Biophys Acta. 2003;1642(1-2):115–23. doi: 10.1016/s0167-4889(03)00105-8. PubMed PMID: 12972300.

49. Uppaluri S, Brangwynne CP. A size threshold governs Caenorhabditis elegans developmental progression. Proc Biol Sci. 2015;282(1813):20151283. doi: 10.1098/rspb.2015.1283. PubMed PMID: 26290076; PubMed Central PMCID: PMCPMC4632629.

50. Rooney JP, Luz AL, Gonzalez-Hunt CP, Bodhicharla R, Ryde IT, Anbalagan C, et al. Effects of 5’-fluoro-2-deoxyuridine on mitochondrial biology in Caenorhabditis elegans. Exp Gerontol. 2014;56:69–76. Epub 20140403. doi: 10.1016/j.exger.2014.03.021. PubMed PMID: 24704715; PubMed Central PMCID: PMCPMC4048797.

